# A systematic analysis of the beta hairpin motif in the Protein Data Bank

**DOI:** 10.1101/2020.10.28.359612

**Authors:** Cory D. DuPai, Bryan W. Davies, Claus O. Wilke

**Affiliations:** Department of Molecular Biosciences, University of Texas at Austin, Austin, Texas, USA; Department of Integrative Biology, University of Texas at Austin, Austin, Texas, USA; Center for Systems and Synthetic Biology, John Ring LaMontagne Center for Infectious Diseases, Institute for Cellular and Molecular Biology, University of Texas at Austin, Austin, Texas, USA

**Keywords:** Beta hairpin, computational biology, PDB, protein design

## Abstract

The beta hairpin motif is a ubiquitous protein structural motif that can be found in molecules across the tree of life. This motif, which is also popular in synthetically designed proteins and peptides, is known for its stability and adaptability to broad functions. Here we systematically probe all 49,000 unique beta hairpin substructures contained within the Protein Data Bank (PDB) to uncover key characteristics correlated with stable beta hairpin structure, including amino acid biases and enriched inter-strand contacts. We also establish a set of broad design principles that can be applied to the generation of libraries encoding proteins or peptides containing beta hairpin structures.

**Importance:** The beta hairpin motif is a common protein structural motif that is known for its stability and varied activity in diverse proteins. Here we use nearly fifty thousand beta hairpin substructures from the Protein Data Bank to systematically analyze and identify key characteristics of the beta hairpin motif. Ultimately, we provide a set of design principles for the generation of synthetic libraries encoding proteins containing beta hairpin structures.

## Introduction

Beta hairpins, one of the simplest stable protein structural elements, consist of two antiparallel beta-sheets joined by a short loop region. Despite their simplicity in form, beta hairpins are highly adaptable in function. Beta strands are known to participate in protein-protein interactions that are often facilitated by specific amino acid orientations^1^ and beta hairpin motifs are no different.^2–4^ Indeed, these motifs are a core feature in a diverse array of bioactive molecules, from large beta barrel proteins that transport cargo through cellular membranes^5–7^ to substantially smaller antimicrobial peptides and peptide derivatives.^8–10^ Whether through self-aggregation,^11,12^ target binding,^13^ or amphipathic structure formation,^6,14^ beta hairpin motifs facilitate a range of different biological functions.

In addition to its prevalence in nature, the beta hairpin motif is stable in even small structures and extensively adaptable to specific functions, making it a popular choice in engineered protein structures. Efforts to design such structures have benefited from several decades of research aimed at identifying how beta hairpins form^15–17^ and what factors influence their stability and specific activity.^2,18–21^ Examples of synthetic proteins that have successfully adapted the beta hairpin motif for specialized functions include hydrogels,^9^ antimicrobial peptides,^22^ and various molecules with material science applications.^8^ Although largely successful, beta hairpin engineering efforts are typically limited to testing relatively small libraries involving derivatives of a stable scaffold structure or existing protein via peptidomimetics.^2,4,19,23,24^ With the increasing availability of high throughput screening platforms to test for activity in large libraries of de novo sequences^25–27^ there is an obvious need for broader design principles that can be applied to the generation of libraries with millions of diverse beta hairpin containing proteins. Knowledge of amino acid propensities throughout known beta hairpin sub-structures could inform such design principles but existing catalogs are too broadly focused on beta sheets, outdated, or limited in scope.^16,20,28–31^ An up-to-date characterization of amino acid distributions at specific positions within beta hairpins does not exist.

Using a systematic analysis of sequence and structural data from all beta hairpin containing proteins in the Protein Data Bank (PDB), we derived key sequence factors and patterns common to beta hairpins. Important features include amphipathic faces created by the periodic alternation of hydrophilic and hydrophobic amino acids within beta strands, the high prevalence of aspartic acid/asparagine caps at the N-terminal end of beta strands, and specific residue contacts that are over (e.g. cysteine-cysteine, salt bridges) and under (e.g. proline-lysine) represented. These findings give us a broader understanding of naturally occurring beta hairpins and will aid future efforts in the design of bioactive molecules containing the beta hairpin motif.

## Results

### General approach

To identify and classify motifs we used the following process (see Materials & Methods for further detail). We first collected all PDB structures^32^ and their corresponding amino acid sequences filtered to 90 percent similarity. We then used DSSP-derived secondary structure annotations^33^ to identify potential beta hairpin substructures consisting of two antiparallel beta-sheets joined by a short loop region (Fig. 1). After determining contacting residues between beta strands, we excluded any structures with less than four contacts from further analysis. This process identified nearly 50,000 unique beta hairpin motifs from some 24,000 independent protein structures. Using these structures, we calculated average amino acid frequencies within structural regions and observed amino acid contacts between hairpin beta strands. We then classified and divided motif structures based on turn length and orientation of beta strand faces. Using these groupings, we determined average amino acid frequencies at each position of the beta hairpin motif.

**Figure 1:**
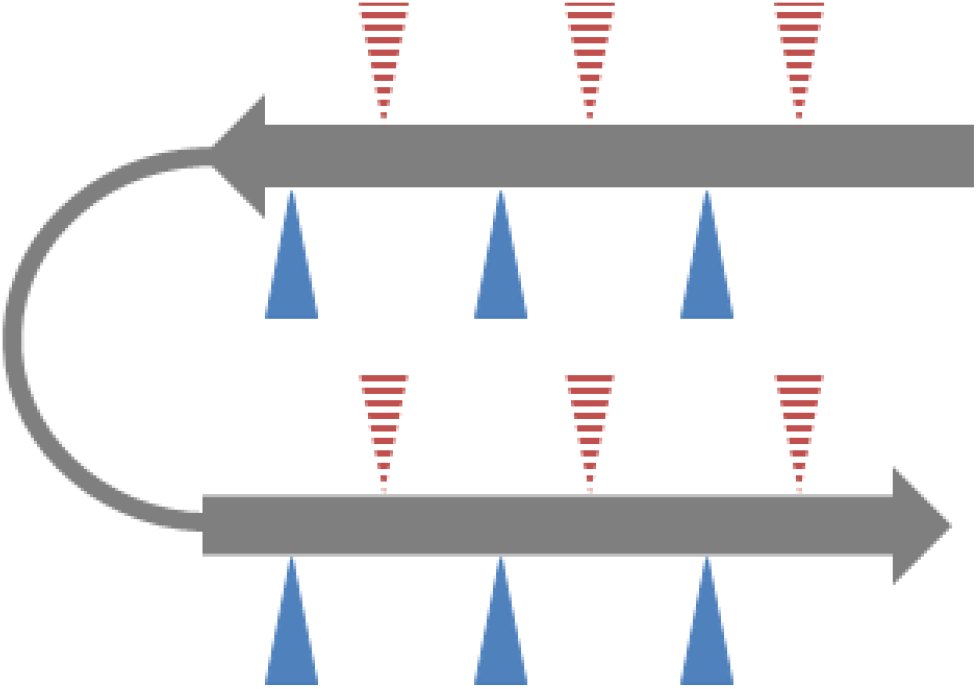
General beta hairpin structure. Beta hairpins consist of two anti-parallel beta strands (grey arrows) linked with a flexible turn region (grey line). Beta strands typically have amphipathic characteristics conferred by alternating hydrophobic and hydrophilic residues. Triangles represent beta strand side amino acid side chains, with red indicating hydrophobic and blue indicating hydrophilic residues. Dashed triangles indicate side chains oriented away from the viewer while solid triangles indicate side chains oriented towards the viewer.

### Secondary structure explains average amino acid frequencies

It has long been known that different secondary structural elements tend to favor the inclusion of certain amino acids over others.^29,30,34,35^ This is exactly what we see with our analysis of beta hairpin motifs (Fig. 2), with a clear difference in average amino acid frequencies between beta strands, the turn region, and background levels across all included protein structures. Our analysis agrees with previous work illustrating a strong preference for glycine, asparagine, and aspartic acid in flexible turn regions.^29,30^ While proline is also more common in the turn region than in either beta strand, we see no difference in turn region prevalence when compared to background levels. This is in contrast to previous findings that saw significant enrichment of proline in turn regions.^8,36^ This lack of proline enrichment and the relatively low average proline abundance in the turn region is particularly surprising given the known role of such residues in stabilizing beta turns.^36,37^

**Figure 2:**
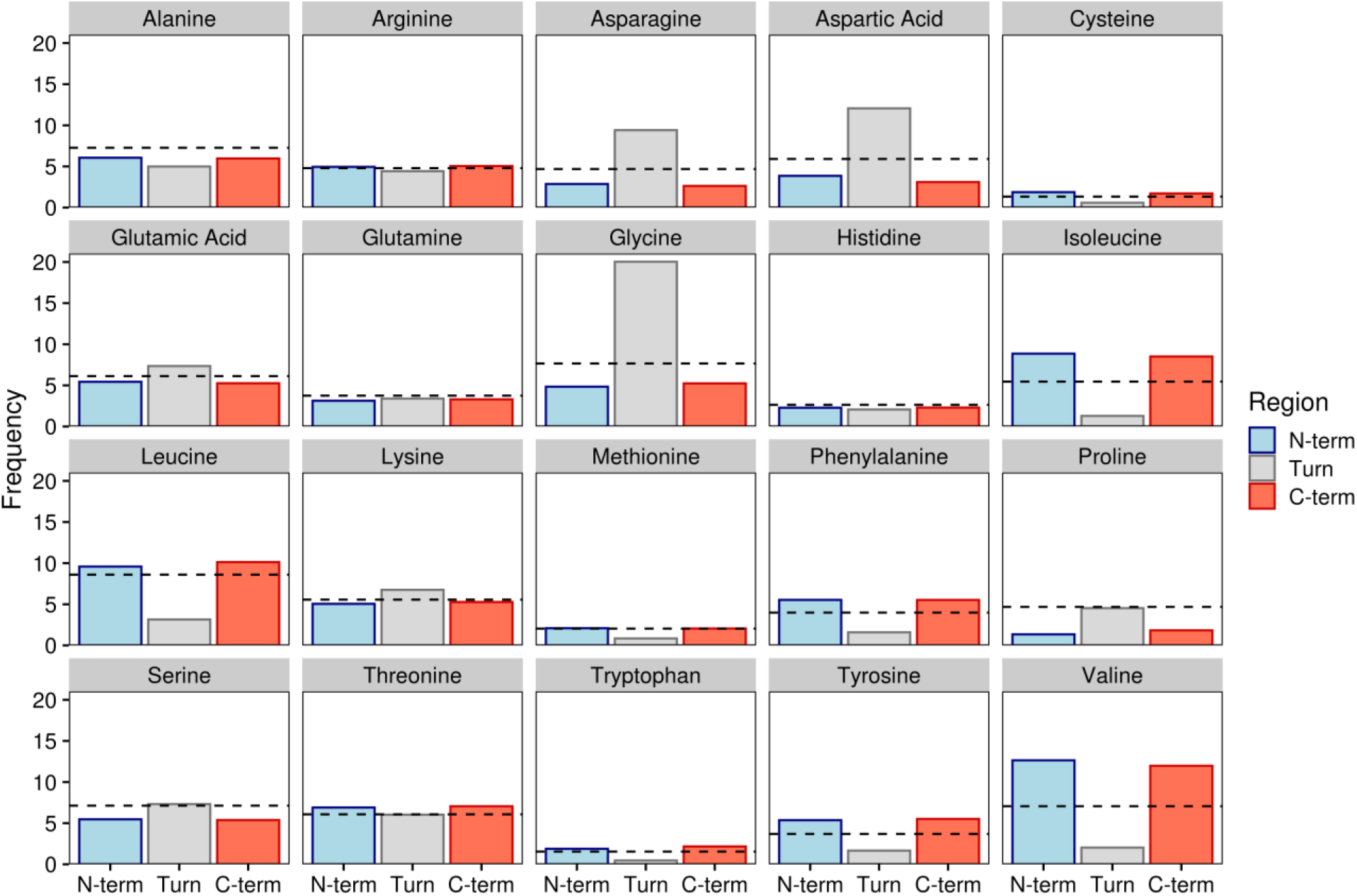
Amino acid frequencies by beta hairpin secondary structure region. Bars indicate average amino acid frequencies for each amino acid within a given region of all beta hairpins. The black dashed line indicates background amino acid frequencies for all sites in all proteins containing the beta hairpin motif. N-term and C-term refer to the N- and C-terminal beta strands while turn denotes the turn region.

When looking at amino acid levels in the beta strands, there appears to be little to no difference in prevalence between strands. Both strands show an increased occurrence of isoleucine, valine, and several other chiefly hydrophobic residues in beta sheet structures, supporting previous research.^38^ Additionally, both strands show a greater tolerance for positively charged residues as is commonly observed with anti-parallel beta strands as opposed to their parallel counterparts.^7,10,39^ We further probed for differences across domains of life but saw no strong trends in individual amino acids (Supp. Fig. 1A). There were, however, taxa specific differences in turn region preference for polar and negatively charged amino acids (Supp. Fig. 1B).

### Residue positional biases are linked to flexibility, stability, and hydrophobicity

Beta hairpins, especially those in membrane interacting structures such as beta barrels and some antimicrobial peptides, are known to incorporate amphipathic beta sheets that periodically alternate between hydrophilic and hydrophobic amino acids, creating two distinct faces^40,41^ (Fig. 1). To account for these faces in our analysis, we divided our dataset based on the presentation of an initial polar or hydrophobic face for both the N and C terminal beta strands (see Materials and Methods). After accounting for these amphipathic faces as well as differences in turn region length, clear patterns emerged in all regions of the beta hairpin motif (Fig. 3). The most obvious pattern observed was the alternating preference for charged/polar and hydrophobic residues in both beta strands (Fig. 3A-B). While hydrophobic residues appear to be more favorable in either beta strand on average (Fig. 2), polar and charged residues are well tolerated when oriented correctly.

**Figure 3:**
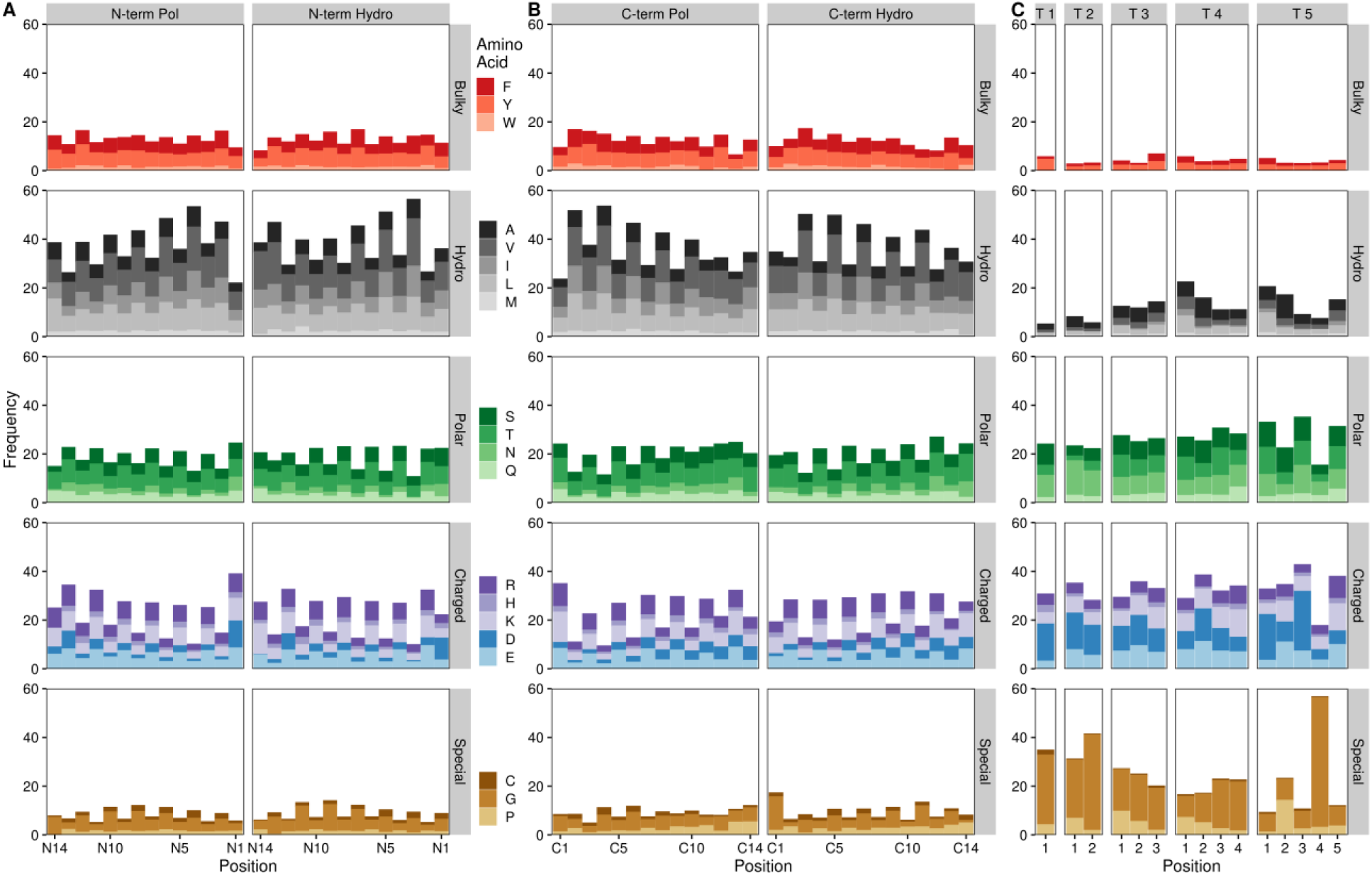
Amino acid frequencies by beta hairpin residue position. Bars indicate average amino acid frequencies for each amino acid at a given position across all beta hairpin structures. N-term and C-term refer to the N- and C-terminal beta strands while T # denotes a turn region of a given length (e.g. T 3 indicates a three residue turn region). Pol refers to beta strands containing a polar face adjacent to the turn region, Hydro denotes a hydrophobic face at this position. Beta strand residues are numbered from the turn region, with residue 1 representing the residue closest to the turn. Turn residues are numbered from N-terminal (residue 1) to C-terminal.

On a more granular level, we further surveyed for differences in amino acid frequencies at specific locations within the larger hairpin motif. In contrast to their average beta strand frequencies, hydrophobic amino acids are also less tolerated at the C-terminal edge of either beta strand regardless of orientation. In their place, aspartic acid and (to a lesser extent) asparagine are over-represented at these loci, with this effect being particularly strong for the N-terminal beta strand where the last residue is one of these two amino acids in nearly 20% of observed hairpins. This frequency is roughly that observed for these two amino acids, on average, in the turn region (Fig. 2, Fig. 3C), although other common turn and cap-associated residues, namely glycine and proline, do not show an over-representation at these positions. Interestingly, aspartic acid residues at the C-terminal end of either beta strand also correlate with increased frequencies of bulky aromatic amino acids (i.e. tyrosine, tryptophan, and phenylalanine) at the N-terminally adjacent position and a preference for glycine at the first N-terminal strand residue (Supp. Fig. 2A).

Although proline showed no enrichment in the average turn region compared to background levels (Fig. 2), proline frequencies are slightly higher than background in the first residue of turns with three to four amino acids and substantially higher than background in the second residue of turns with five amino acids (Fig. 3C). These findings largely agree with existing evidence on the prevalence and importance of prolines in the beginning of turn regions^42–44^ but the nearly four-fold enrichment for residue two prolines in hairpin structures with five amino acid long turn regions when compared to background levels is particularly surprising. In combination with the fact that over half of all fourth residues in five amino acid long turn regions are glycines, these findings suggest that beta hairpins with longer turn regions may have very specific physiochemical requirements that limit amino acid diversity.

### Amino acid contacts between strands favor stabilizing interactions

As the overall beta hairpin structure is stabilized by interactions between the two beta strands, we sought to identify enriched amino acid pairings between strands to see if certain interactions were more common than expected. Pairings between residues with similar electrostatic properties, that is two hydrophobic residues or a polar residue and a polar/charged residue, were largely more common than expected (Fig. 4, Supp. Fig. 3). This data agrees with our previous findings regarding the grouping amino acids into beta strand faces based on similar physiochemical properties. In a similar vein to the pairing of electrochemically similar residues, oppositely charged residues tended to pair together in electrostatically favorable salt bridges that are known to stabilize protein structures.^45–48^ Such salt bridges represented some of the most enriched amino acid pairings.

**Figure 4:**
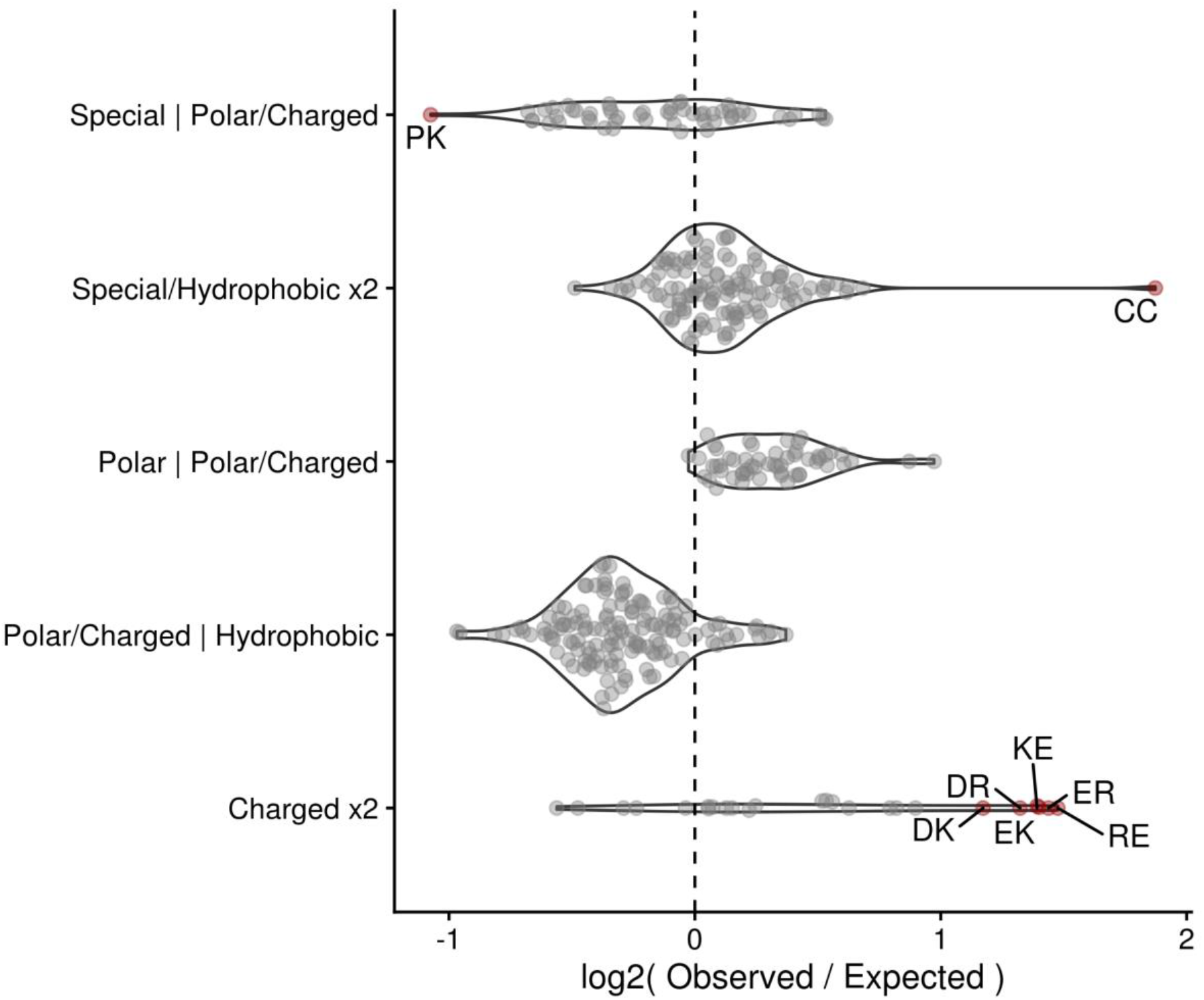
Grouped differences in observed vs. expected residue contacts. Dots represent individual contacting pairs with red, labelled dots indicating contacts that are enriched or depleted at least two-fold vs. expected values. Residues are grouped as follows: Special refers to cysteine, proline, glycine; Hydrophobic refers to valine, leucine, isoleucine, methionine, alanine, tryptophan, tyrosine, phenylalanine; Polar refers to glutamine, threonine, serine, asparagine; Charged refers to arginine, histidine, lysine, aspartic acid, and glutamic acid.

The most enriched amino acid pairing between beta strands is that of cysteine with itself to create a structurally stabilizing di-sulfide bond. Such pairings are often used to stabilize engineered peptide structures^49,50^ and cysteine coupling is so preferential in nature that many organisms possess a proteome-wide bias towards even numbers of cysteine residues.^51^

In contrast to enriched contact pairings, several classes of interactions, typically those between electrochemically dissimilar residues, were observed much less than expected. The low observance of inter-strand contacts between polar/charged and hydrophobic amino acids (Fig. 4) is intuitive given the strong repulsive nature between such residues which could destabilize overall protein structure.

### Design Principles

Taken altogether, our work provides a strong foundation of general principles that can be applied to the design of functionally diverse high throughput beta hairpin libraries (Table 1). First, libraries should seek to incorporate beta strands with amphipathic faces as seen in our analysis of beta strand positional biases (Fig. 3A-B). Second, aspartic acid and asparagine should be favored at C-terminal beta strand residues, especially in the beta strand preceding the turn region. Next, proline and glycine should be utilized in residues two and four of five residue turn regions given their overwhelming enrichment in these positions (Fig. 3C). Fourth, average secondary-structure amino acid preferences should inform design choices, especially within the turn region. While residues in both hairpin beta strands show positionally specific frequency deviations from secondary structure averages (Fig. 2, Fig. 3A-B), there is much less deviation within the turn region (Fig. 3C). Lastly, stabilizing interactions should be favored between beta strands. Such interactions include salt bridges, disulfide bonds, and the pairing of certain biochemically similar residues (i.e. hydrophobic-hydrophobic and polar-polar pairings) (Fig. 4). These simple guidelines are specific enough to inform design choices while flexible enough to allow for applications across broad research areas.

**Table 1:** Design principles

| Table 1: Design principles |
| --- |
| 1. Incorporate amphipathic beta strand faces |
| 2. Favor aspartic acid/asparagine at C-terminal beta strand residues |
| 3. Incorporate T2 proline and T4 glycine in five residue turns |
| 4. Account for secondary structure biases, especially in the turn region |
| 5. Favor salt bridges and di-cysteine interactions to provide stability |

## Discussion

By analyzing the composition of beta hairpin motifs across all proteins within the PDB we have identified key characteristics of this versatile structure. Expanding on existing knowledge of secondary structure biases, we outline the preference for the amphipathic orientation of amino acids within beta strands to create two faces with different physiochemical properties. We further identify key positional preferences for specific amino acids in all regions of the hairpin motif. Lastly, we highlight the importance of stabilizing interactions between residues in the N and C terminal beta strands of the hairpin.

Our results integrate and expand upon existing knowledge of protein amino acid biases and intra-protein interactions to provide a systematic framework and novel insights to describe the beta hairpin motif. We find that stable beta hairpin structures tend to possess site-specific amino acid preferences and to incorporate amphipathic character in both hairpin beta strands. While-existing secondary-structure-specific amino acid distributions ^29,30^ are accurate and informative, such averages prove inadequate to capture the inherent nuances of the beta hairpin motif. For instance, while our analysis finds that an average hairpin beta strand would consist of only hydrophobic residues (Fig. 2), a beta hairpin containing two such average strands without any amphipathic character would be statistically improbable (Fig. 3A-B) and highly unlikely to fold correctly ^21^, let alone function biologically ^10^.

Position-specific amino acid biases need to be considered to help form stable protein structures. Our observation that prolines are less enriched in turn regions (Fig. 2) than previously observed ^8,36^ is perhaps best explained by the extreme position-specific preference of proline residues in turn regions of a given length (Fig. 3C). Thus, certain proline residues are enriched within and likely to stabilize hairpin turn regions even though there is no strong trend when averaged across all turn residues. Outside of the turn region, hairpin beta strands also exhibit amino acid biases at key loci as well as a strong proclivity to incorporate stabilizing inter-strand contacts. We find that asparagine and aspartic acid residues are much more common at the C-terminal end of either hairpin beta strand (Fig. 3A-B, Supp. Fig. 2). These residues may participate in a beta capping phenomenon to block the continuation of beta structure into a turn region ^16^. A beta capping role may also explain our observation of an increased prevalence of bulky aromatic residues preceding terminal aspartic acids (Supp. Fig. 2) as aromatic residues are known to stabilize beta hairpin structures ^18,19^. Lastly, appropriate contacts between hairpin beta strands are imperative to provide structural stability. As an example, we identified cysteine pairings as being particularly enriched in beta hairpin substructures (Fig. 4). Such pairings have long been used to stabilize engineered peptide structures ^49,50^, are so preferential in nature that many organisms possess a proteome-wide bias towards even numbers of cysteines ^51^.

While our analysis of amino acid preferences within beta hairpin secondary structures across the domains of life showed no strong differences (Supp. Fig. 1A) there were some interesting minor trends as well as a notable difference in turn region composition between taxa (Supp. Fig. 1B). Cysteines, which are fairly uncommon across proteins in general, appear twice as often in Eukaryotic beta hairpins than in Prokaryotic or Archaeaotic beta hairpins. This observation agrees with previous data showing the same trend of increasing cysteine occurrence in proteomes of more complex organisms ^52–55^. Of greater note is the inverse relationship between polar and negative amino acid propensities within beta hairpin turn regions across taxa. Frequencies for negatively charged amino acids within the turn region decrease from Archaea to Bacteria, Eukarya, and finally Viruses while polar amino acids show the opposite trend. This difference is likely explained by protein adaptations to harsh environments in Archaea/Bacteria ^56^ that are less commonly encountered by Eukaryotic or viral proteins. This trend is not seen in either beta strand of the hairpin as turn structures are some of the most accessible protein regions ^57^ and would likely experience more selective pressure in harsh environments than less exposed beta strands.

One major limitation of our approach is that we were only able to establish broad general properties of beta hairpins that might influence overall structure or function. This is in contrast to prior work that has focused on identifying key design factors for specific beta hairpin scaffolds ^23,24,42,58^ or grouping beta hairpins and related structures into increasingly detailed classifications ^20,31,44^. While the PDB dataset that we analyzed could be used to expand upon these highly focused areas of research, the broad applicability of our results would be compromised.

In combination with prior research efforts, our simple design guidelines (Table 1) can be adapted to the creation of large-scale protein or peptide libraries aimed at almost any functional purpose, from anticancer drugs to biosensors. For example, beta hairpin antimicrobial peptides are known to incorporate multiple disulfide bonds and favor an overall net positive charge while still maintaining amphipathic character ^10,13^. Adapting our design principles with these properties in mind would facilitate the construction of a library of positively charged, disulfide stabilized peptides with presumptive beta hairpin structure to test for antimicrobial activity.

In summary, our findings are broadly adaptable to creating large libraries of beta hairpin containing molecules skewed towards a specific functionality and will help engineering efforts keep pace with the ever-expanding capacity of screening assays.

## Materials and Methods

### Identification of beta hairpin substructures

We defined the beta hairpin motif as an amino acid sequence containing two sets of four to fourteen extended beta strand residues joined by one to five turn, bend, or unannotated residues. A maximum beta strand length of fourteen was selected based on the typical length of beta strands in monomeric beta barrel proteins ^59^ while the range of turn lengths was selected based on prior research into beta hairpins ^17^. We searched DSSP ^33^ derived secondary structure annotations of all PDB proteins (downloaded from https://cdn.rcsb.org/etl/kabschSander/ss.txt.gz on July 22^nd^ 2020) for this motif. We further filtered our dataset to include only IDs for representative structures clustered to within 90% sequence identity. Clusters were obtained from PDB on July 22^nd^ 2020 using the RESTful Web Service Interface (https://www.rcsb.org/pdb/software/rest.do). Further manual filtering was applied to exclude redundant and overly similar hairpin sequences, largely from structures of nanobodies, antibodies, and their derivatives.

### Identification of contacting residues

To ensure that our analyzed motifs possessed the correct beta hairpin 3D structure, we filtered our dataset to only include structures in which at least four amino acid side chain pairs formed contacts between the N and C terminal beta strands. We defined contacts as any pair of residues in which side-chain beta carbons were within 8 Angstroms of one another. Determining contacts via the presence of backbone hydrogen bonds produced similar results (data not included). To calculate expected contact frequencies, individual amino acid frequencies were derived using the relative occurrence of each amino acid across all contact pairs. Values for amino acids in a pairing were then multiplied together to establish an expected frequency for every possible pairing of amino acids.

### Grouping of beta hairpin substructures

To characterize the amphipathic faces of each beta strand, solvent accessibility was averaged across odd and even numbered amino acid residues with the first amino acid being the residue closest to the turn region. Strands in which the odd amino acid residues have a higher mean accessibility were categorized as polar while strands with the opposite phenotype were categorized as hydrophobic. Solvent accessibility was chosen in lieu of hydrophobicity or other metrics as PDB structures contain accessibility information and solvent accessibility is known to correlate with hydrophobicity ^57^.

### Data and figures

All data was analyzed in R using the tidyverse family of packages ^60^ in combination with the data.table ^61^ and seqinr ^62^ packages. All figures were created using ggplot2 ^63^ and cowplot ^64^. Supplementary Figure 3 additionally utilized the ggseqlogo package ^65^. All processed data and analysis scripts are available at https://doi.org/10.5281/zenodo.4069580.

---

**Supplementary Figure 1:**
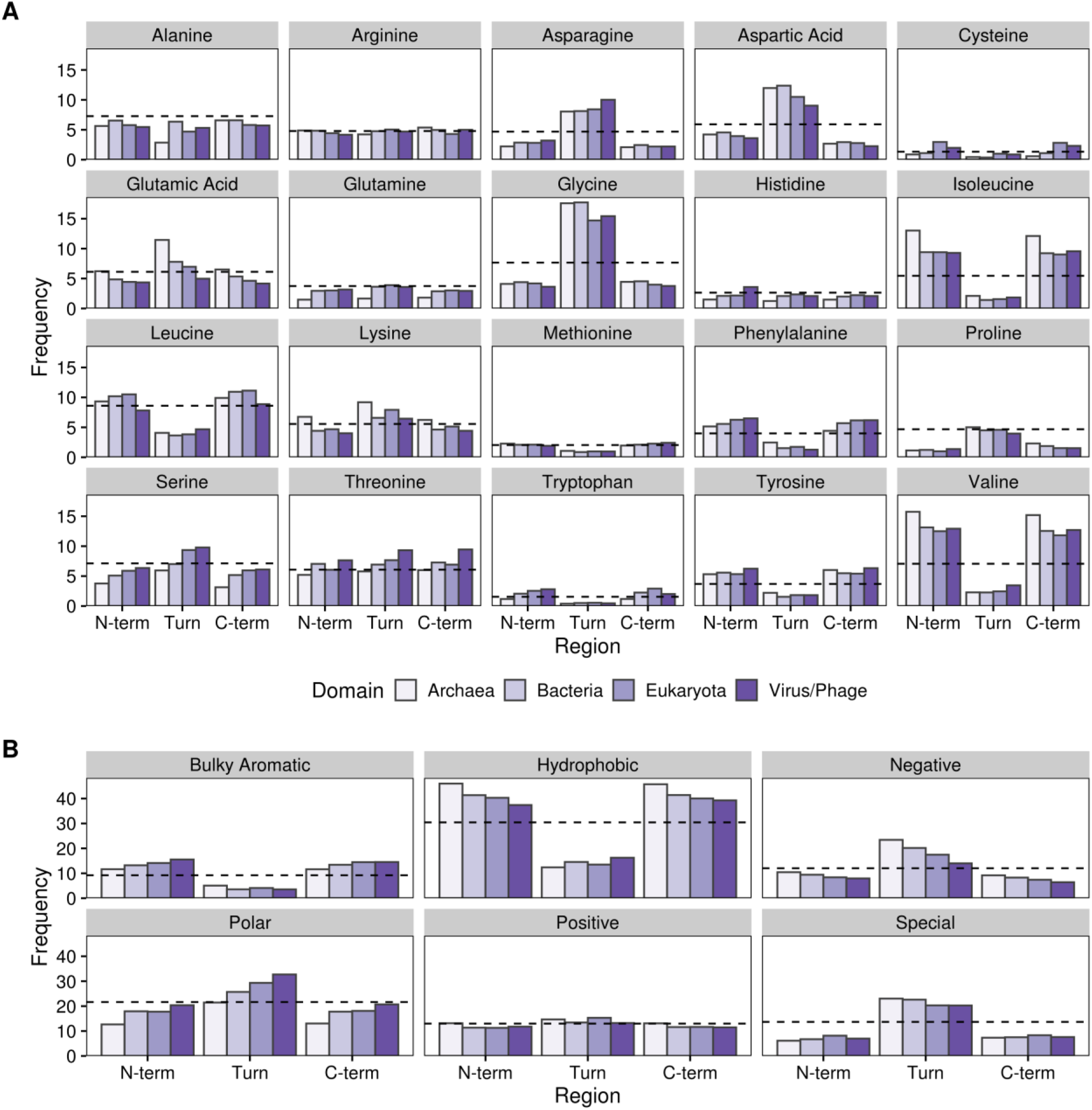
Amino acid frequencies by beta hairpin secondary structure region and organism domain. A) Bars indicate average amino acid frequencies for each amino acid within a given region of all beta hairpins broken down by Domain of origin. The black dashed line indicates background amino acid frequencies for all sites in all proteins containing the beta hairpin motif. N-term and C-term refer to the N- and C-terminal beta strands while turn denotes the turn region. B) Same as in A) with amino acid frequencies grouped into specific classes. Amino acids are grouped as follows: Bulky Aromatic - phenylalanine, tryptophan, tyrosine; Hydrophobic - alanine, isoleucine, leucine, methionine, valine; Negative - aspartic acid, glutamic acid; Polar - asparagine, glutamine, serine, threonine; Positive - arginine, histidine, lysine; Special - cysteine, glycine, proline.

**Supplementary Figure 2:**
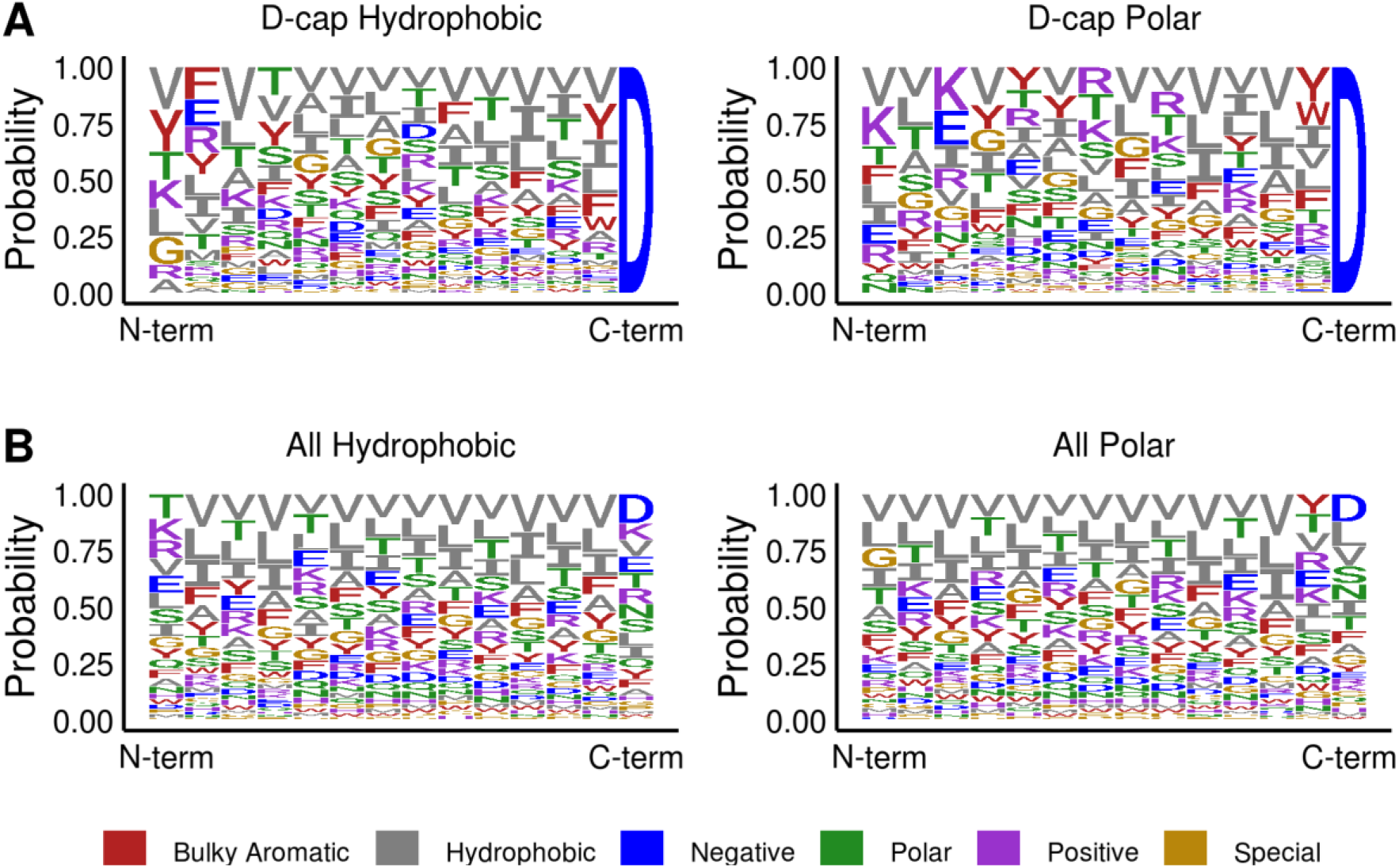
Motif logo of amino acid probabilities in hairpin beta strands. Amino acid letters are scaled to represent relative frequency at a given position in both beta strands, with more common amino acids listed first vertically. Color indicates the type of amino acid. A). Logo motif for hairpin beta strands with a C-terminal aspartic acid that start with either a hydrophobic (left) or polar (right) face relative to the turn region. B). As in A) but showing frequencies across all hairpin beta strands, not just those terminated with an aspartic acid residue. Amino acids are grouped as in Supp. Fig. 1.

**Supplementary Figure 3:**
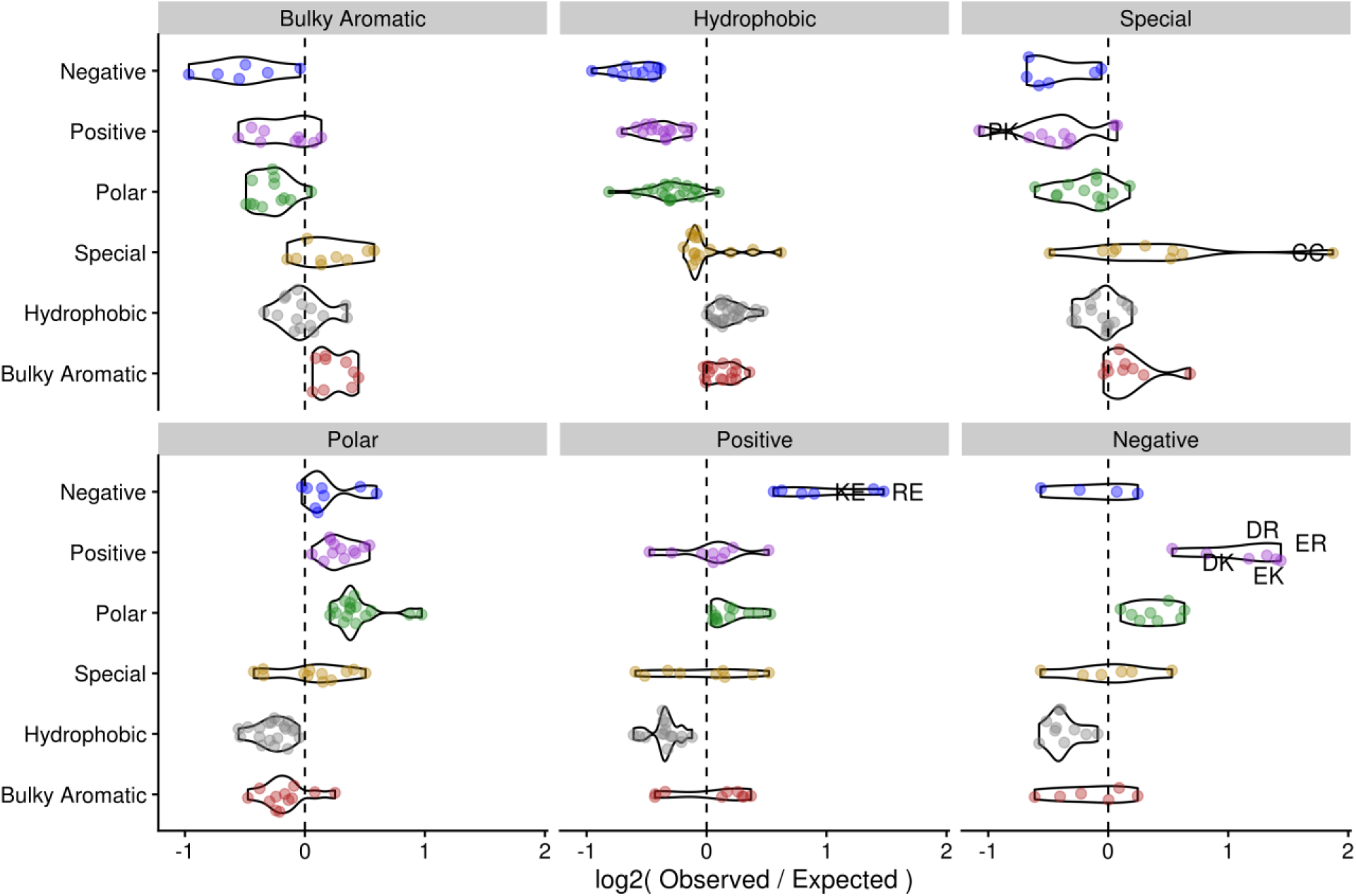
Differences in observed vs. expected residue contacts. Dots represent individual contacting pairs with labelled dots indicating contacts that are enriched or depleted at least two-fold vs. expected values. Y-axis labels denote the class of the first residue in the pair while facet headings denote the class of the second amino acid (e.g. the dot for DP would be in the Negative row of the Special column). Amino acids are grouped as in Supp. Fig. 1.

